# Ancient farmer and steppe pastoralist-related founding lineages contributed to the complex landscape of episodes in the diversification of Chinese paternal lineages

**DOI:** 10.1101/2023.08.28.555114

**Authors:** Mengge Wang, Yuguo Huang, Kaijun Liu, Haibing Yuan, Shuhan Duan, Zhiyong Wang, Lanhai Wei, Hongbing Yao, Qiuxia Sun, Jie Zhong, Renkuan Tang, Jing Chen, Yuntao Sun, Xiangping Li, Haoran Su, Qingxin Yang, Liping Hu, Libing Yun, Junbao Yang, Shengjie Nie, Yan Cai, Jiangwei Yan, Kun Zhou, 10K_CPGDP Consortium, Chuanchao Wang, Bofeng Zhu, Chao Liu, Guanglin He

## Abstract

Ancient DNA advances have reported the complex genetic history of Eurasians, but how the knowledge of ancient subsistence strategy shifts and population movements influenced the fine-scale paternal genetic structure in East Asia has not been assessed. Here, we reported one integrated Y-chromosome genomic database of 15,530 people, including 1753 ancient people and newly-reported 919 individuals genotyped using our recently-developed targeted sequencing YHSeqY3000 panel, to explore Chinese genomic diversity, population evolutionary tracts and their genetic formation mechanism. We identified four major ancient technological innovations and population movements that shaped the landscape of Chinese paternal lineages. First, the expansion of millet farmers and early East Asians from the Yellow River Basin carrying the major O2/D subclades promoted the formation of the Sino-Tibetan people’s major composition and accelerated the Tibetan Plateau’s permanent occupation. Second, rice farmers’ dispersal from the Yangtze River Valley carrying O1 and some sublineages of O2 contributed significantly to Tai-Kadai, Austronesian, Hmong-Mien, Austroasiatic people and southern Han Chinese. Third, Siberian-related paternal lineages of Q and C originated and boomed from Neolithic hunter-gatherers from the Mongolian Plateau and the Amur River Basin and significantly influenced the gene pools of northern Chinese. Fourth, western Eurasian-derived J, G and R lineages initially spread with Yamnaya steppe pastoralists and other proto-Indo-European people and further widely dispersed via the trans-Eurasian cultural communication along the Eurasian Steppe and the ancient Silk Road, remaining genetic trajectories in northwestern Chinese. Our work provided comprehensive modern and ancient genetic evidence to illuminate the impact of population interaction from the ancient farmer or herder-based societies on the genetic diversity patterns of modern people, revised our understandings of ancestral sources of Chinese paternal lineages, underscored the scientific imperative of the large-scale genomic resources of dense spatiotemporal underrepresented sampling populations to understand human evolutionary history.

## Introduction

Comprehensively documenting the genetic landscape of genetically diverse worldwide populations and illuminating their influence on the human demographical history and genetic basis of complex traits and diseases was the goal of population genomics and human pangenome projects (Bergstrom et al. 2020; Byrska-Bishop et al. 2022). Current genetic resources, especially large-scale population genomic cohorts, were mainly derived from descendants of European ancestry, which limited the transferability of European-based genetic findings in other non-European people as their differences in the allele frequency spectrum, linkage disequilibrium, effect size and different trajectories of the evolutionary process (Sirugo et al. 2019). Previous anthropological and genetic findings suggested that the genetically underrepresented African continent possessing the most linguistic (Afro-Asiatic, Nilo-Saharan, Niger-Congo and KhoeSan) and genetic diversity was regarded as the cradle of modern human origin (Choudhury et al. 2020). Similarly, East Asia, including countries of China, Mongolia, Japan, North Korea and South Korea, served as one of the earliest cradles of civilization and the crossroad of the peopling of Oceania, Siberia and America, whose genetic landscape is also poorly characterized in the era of population-based genomics. China is the world’s second-most populous country with a population size exceeding 1.4 billion, and there are nearly 300 living languages that belong to seven language families [Sino-Tibetan (ST), Altaic (Mongolic, Tungusic, Turkic, Japonic and Koreanic), Tai-Kadai (TK), Hmong-Mien (HM), Austronesian (AN), Austroasiatic (AA) and Indo-European (IE)] (Lewis et al. 2016). China harbors substantial genetic, physical, cultural and ethnolinguistic diversity, which allows this region to maintain an unrivaled position in the study of the complex demographic history of ethnolinguistically diverse populations, including human divergence, migration and admixture events, as well as interrelationships between genetics and cultures (Wang, Yeh, et al. 2021; Kumar et al. 2022; Zhang et al. 2022). It has been well-documented that European bias can cause human health inequality and non-transferability of PRS across genetically different populations (Sirugo et al. 2019). The worldwide genetic consortiums have conducted many studies or projects to fill the gap in the missing genetic diversity in human genetics and genomics and dissect the genetic basis of complex traits/diseases and the evolutionary history of Chinese populations. Recently, a large body of research focused on geographically different ST, Mongolic, Tungusic, Turkic, TK and HM groups has been carried out using genome-wide SNP arrays (Feng et al. 2017; He, Wang, Li, et al. 2021; Ma et al. 2021; He et al. 2022; Wang, He, et al. 2022; Wang et al. 2023). Moreover, there has been a rapid increase of whole-genome sequencing (WGS) studies on ethnolinguistically diverse Chinese populations, such as the Westlake BioBank for Chinese (WBBC), NyuWa genome resource, China Metabolic Analytics Project (ChinaMAP), 10K Chinese People Genomic Diversity Project (10K_CPGDP) and STROMICS (Cao et al. 2020; Zhang, Luo, et al. 2021; Cong et al. 2022; Cheng et al. 2023; He, Yao, et al. 2023). Generally, these studies advanced our understanding of demographic history, the patterns of genetic diversity and the genetic architecture of complex traits/diseases in Chinese populations from autosome-related perspectives. More unknown genetic features from a uniparental perspective and population-scale project should be further explored.

A complex assortment of neutral and non-neutral selection processes has shaped the genetic landscape of present-day humans. Autosome-based genomic studies could provide basal insights into the population demographic and adaptive history, and mitochondrial and Y-chromosomal variants could provide additional signals of evolutionary processes that have uniquely left signatures in male/female specific regions in the Y/mitochondrial DNA (mtDNA) genome (Poznik et al. 2016; Nielsen et al. 2017; Li et al. 2019). The non-recombining part of the Y-chromosome (NRY) is strictly paternally inherited, transmitted through the germ line and not affected by heteroplasmy (Jobling and Tyler-Smith 2017), namely unique features of haploidy, escape from crossing over and male specificity. The length of the Y-chromosome is approximately 4,000 times larger than that of mtDNA. Therefore, it contains much more information content relative to mtDNA. The mutation rates of Y-chromosomal single nucleotide polymorphisms (Y-SNPs) are lower than that of mtDNA and short tandem repeats (STRs), and Y-chromosomal variants show more obvious geographical specificity than other markers. These properties have led to its genetic variations becoming a significant part of studies of human evolutionary history on different time scales (Poznik et al. 2016; Jobling and Tyler-Smith 2017).

Improved sequencing technologies and computational innovations in genome assembly, read mapping, variant calling and benchmarking have promoted the generation of many complete Y-chromosomal sequences and greatly enriched our understanding of NRY variations over the past few years (Olson et al. 2023). These identified Y-chromosomal variants enable a robust phylogenetic tree to be gradually constructed, in which the branch lengths are proportional to the number of mutations (Poznik et al. 2016; Jobling and Tyler-Smith 2017; Zhabagin et al. 2022). In addition to constructing a high-quality and high-resolution phylogeny with a robust topology structure and divergence models, several broad approaches have been applied to estimate the mutation rates of Y-SNPs, which could facilitate the conversion of branch length information into time and enable the estimation of the time to the most recent common ancestor (TMRCA) shared by two individuals (Xue et al. 2009; Francalacci et al. 2013; Poznik et al. 2013; Helgason et al. 2015; Jobling and Tyler-Smith 2017).

Many vital genetic works have been conducted to trace one’s ancestor through the paternal lineages and have been an essential source of phylogenetic information for studies of human origin, migration and admixture history in the past two decades of the human genome era (Jobling and Tyler-Smith 2017). Su et al. genotyped 19 Y-SNPs in 925 geographically and genetically different male samples and identified a tremendous northward migration into East Asia during the last Ice Age (Su et al. 1999). Soon after, Ke et al. genotyped three Y-chromosomal biallelic markers in 12,127 male individuals from 163 populations and confirmed the Out-of-Africa hypothesis (Ke et al. 2001). Y-chromosomes can also document trajectories of recent population migration, admixture and expansion events. Zerjal et al. investigated the genetic diversity of 2,123 males using genotype data of 32 Y-chromosomal markers, and the observed pattern of the star-cluster phylogeny indicated the significant influence of the Mongol Empire’s westward expansion on the genetic architecture of Asian populations (Zerjal et al. 2003). Large-scale genetic studies focused on the fine-scale paternal demographic histories of East Asians have been conducted over the past two decades, revealing the patterns of admixture and microevolution during the initial human settlement and subsequent migrations in East Asia (Wang and Li 2013). Resequencing whole Y-chromosomes based on next-generation sequencing (NGS) and computational technology has revolutionized the research paradigm. Wei et al. identified 6,662 high-confidence variants in 36 diverse Y-chromosomes and calibrated previously available Y-chromosomal phylogenies (Wei et al. 2013). Poznik et al. reported 1,244 complete Y-chromosomal genomes randomly sampled from 26 worldwide populations by the 1000 Genomes Project (1KGP), and they discovered more than 65,000 variants and found bursts of expansion within specific paternal lineages occurred in the last few thousand years (Poznik et al. 2016). Several Y-chromosomal investigations have also been performed on a single population or specific lineage. Wang et al. analyzed 285 Y-chromosomal sequences and identified two Neolithic expansions of Tibeto-Burman (TB) groups and their specific paternal lineages (Wang et al. 2018). The O1a-M119 shared by Sinitic, TK and AN people, the founding paternal lineages of Tungusic or Mongolic groups (C2a-F5484 or C2b1a1a1a-M407), C2b-F1067 dominated in eastern Eurasian populations, Q1a1a-M120 connecting East Asian and Siberian populations and other specific lineages have been studied to explore their origin, diffusion and contribution to the gene pool of ethnolinguistically diverse East Asian groups (Huang, Wei, et al. 2018; Sun et al. 2019; Wu et al. 2020; Liu, Ma, et al. 2021; Sun et al. 2021). However, the large-scale genomic database was limited for China, which can provide one vital clue for exploring the entire genetic landscape of Chinese people and their ancient influence factors.

To promote the effective screening of East Asian-specific phylogenetically informative markers and develop a high-resolution Y-SNP panel for large-scale population genotyping of ethnolinguistically diverse groups, we initiated the 10K_CPGDP, one of the aims of which is to comprehensively capture the entire genetic landscape of genetically and ethnolinguistically diverse Chinese populations, including the elucidation of Y-chromosomal genetic variations and the paternal demographic history of underrepresented ethnic groups included in the YanHuang Cohort (YHC) genomic resource (He, Yao, et al. 2023). The YHC in the 10K_CPGDP was focused on presenting one high-quality population-specific Y-chromosome database, drawing one time-stamped higher-resolution Y-phylogeny and developing multiple NGS panels for medical and forensic applications via the combination of WGS and third-generation sequencing (TGS) techniques, including SNPs, STRs, InDels and other variations. We have developed the highest resolution chrY-specific targeted resequencing panel, the ’YHSeqY3000’, and designed the SNP composition based on the whole-genome sequences and genome-wide SNP variations of Y-chromosomes in the YHC genomic resource. We genotyped 3002 panel-related Y-SNPs in 919 male individuals from 57 ethnolinguistically diverse Chinese populations and reported the cohort design and fine-scale paternal evolutionary history of Chinese minority ethnic groups. We presented one integrated Y-chromosome database including 15,530 individuals from modern and ancient Eurasian populations to eliminate the impact of ancient population migration, admixture and agricultural innovations on the landscape of the genetic structure of East Asians. We presented the haplogroup frequency spectrum (HFS) of modern and ancient Eurasians and identified multiple steppe pastoralist-related and agriculture-related founding lineages that formed the mosaic diversity of ethnolinguistically diverse Chinese people. The YHSeqY3000 can serve as a unique tool in the interdisciplinary research of evolutionary, population, medical and forensic genetics, such as paternal demographic history reconstruction, patrilineal biogeographic ancestry inference and forensic pedigree searching.

## Results and Discussion

Genetic diversity of YanHuang cohort paternal lineages inferred from the highest resolution chrY-specific targeted resequencing YHSeqY3000 panel YanHuang Cohort genomic dataset (**Fig. 1A**) in the 10K_CPGDP focused on the genetic diversity of male-specific regions of the Y-chromosome, which was the best resource to gain new insights into the full landscape of the paternal evolutionary history of East Asia. To characterize the genetic patterns of paternal lineages of ethnolinguistically distinct Chinese populations at a fine scale, we developed the highest resolution YHSeqY3000 panel based on the newly-updated database of Y-chromosomal sequences from the 10K_CPGDP genomic resource, including newly-identified Y-SNPs not presented in the latest ISOGG Y-DNA and Yfull phylogenetic trees (submitted), and sequenced 919 participants from 57 populations of 39 ethnic minorities using our newly-developed panel (**Fig. 1A** and Table S1). The YHSeqY3000 panel we presented here allowed simultaneous genotyping of more than 3000 Y-SNPs, covering the overwhelming majority of subclades of dominant paternal lineages of Chinese populations. We adopted three haplogroup classification methods, including Y-LineageTracker and HaploGrouper based on ISOGG2019-tree and in-house script based on the newly-reconstructed phylogenetic tree, to perform haplogroup inference simultaneously for reliable classification results. There were forty discrepancies (classified into entirely different macro-haplogroups) between the haplogroup results obtained through Y-LineageTracker and our in-house scripts, mainly including subclades of C-M130, J-M304, N-M231, O-M175, Q-M242, R-M207 and T-M184. However, there were only four discrepancies between the haplogroup results obtained through HaploGrouper and our in-house script; the samples belonging to F-M89 were classified into CF by HaploGrouper. The fundamental analysis for the research on paternal demographic history is the NRY haplogroup inference. Several tools have been developed to support the function of haplogroup classification: Y-LineageTracker (Chen et al. 2021), HaploGrouper (Jagadeesan et al. 2021), Yleaf (Ralf et al. 2018), AMY-tree (Van Geystelen et al. 2013) and cleantree (Ralf et al. 2015), but the NRY haplogroups assigned by some tools were outdated without reference to the latest phylogenetic tree. With the accumulation of Y-chromosome sequencing data, more and more novel NRY variants were identified. However, they have not been located in the up-to-date phylogenetic trees, which also led to errors or inaccuracies in haplogroup inference based on the non-updated phylogenetic topologies. Hence, it is necessary to calibrate and refine the Y-chromosome phylogenetic trees based on newly-identified Y-SNPs and build the continuously updated phylogeny into haplogroup classification tools. The new version of phylogenetic topology to be reported in the 10K_CPGDP will fill this gap. We observed 564 distinct paternal lineages that mainly fell into haplogroups C-M130, N-M231, O-M175 and R-M207, respectively, sampled from 150, 65, 583 and 46 individuals (**Figs. 1B-C** and **S1, Table S2**)). We found that 384 subhaplogroups belonging to C-M130, D-CTS3946, G-M201, J-M304, N-M231, O-M175, Q-M242 and R-M207 were observed only once (**Figs. 1C** and **S1, Table S2**). Significantly, we observed that subclade E1b1b1a-L539 of E-M96, which was found at high frequencies in East Africa, was present in eight individuals from Mongolian, Manchu and Hui populations (**Fig. 1C**). Haplogroup H-L901, one of the most dominant paternal lineages amongst populations in South Asia, was observed in ethnic minority groups in Northwest China. In addition, West Asian-derived T-M184 was observed in Turkic-speaking populations. The patterns of lineage distribution from one geographical region showed that different founding populations contributed to the Chinese paternal gene pool, mainly possibly from ancient migrations from Chinese indigenous rice or millet farmers or western Eurasian steppe people (**Fig. 1B**). We found that haplotype diversity (HD) values reached 1 in all populations with a sample size larger than 30, and haplogroup diversity (H) values ranged from 0.9537 (Zhuang) to 0.9979 (Manchu). Our observations demonstrated that the resolution and coverage of the newly-designed YHSeqY3000 panel are currently the highest, and this panel can be applied for finer haplogroup classification of Chinese populations than previously developed systems (Wang et al. 2019; Liu et al. 2022; He, Wang, et al. 2023; Tao et al. 2023).

**Fig. 1.**
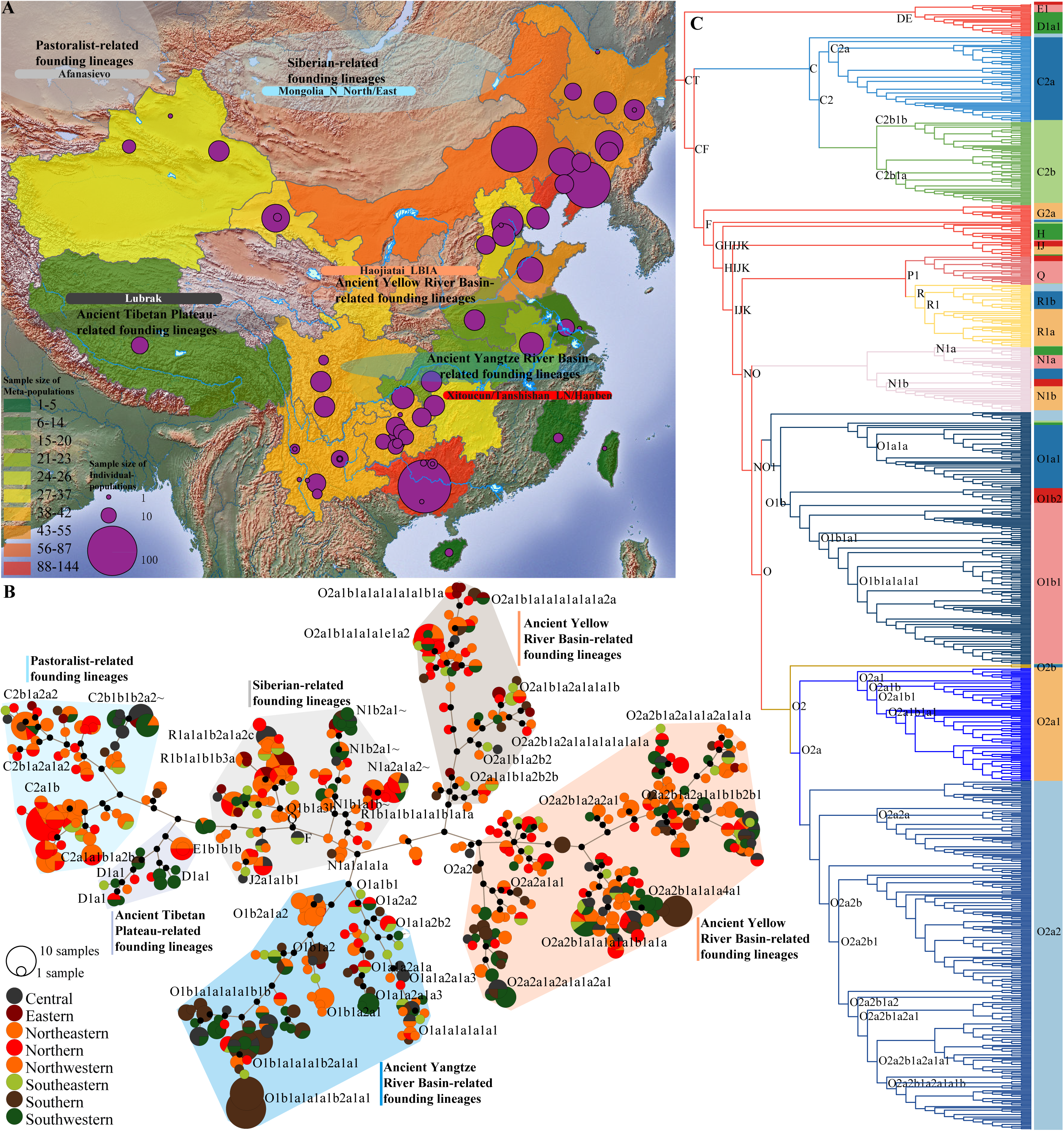
Geographical position, sample size and phylogenetic features of newly-generated 919 targeted sequences. **(A)** A map of East Asia showed the basic information of 919 individuals from 57 geographically or ethnically different populations. The circle size in the map denoted the sample size of individual populations. The colored provinces indicated sampling places, and the color of the province showed the total sample size from these geographical regions. Ancient subsistence strategies (pastoralists, fisher-hunter-gatherers and farmers) from western Eurasia, the Mongolian Plateau and Chinese agriculture original centers (Yellow and Yangtze River Basin) were also denoted. **(B)** Network relationships among 919 haplotypes were inferred based on the median-joining network algorithms. Different colors of the circle showed the geographical origin of one focused haplogroup, and different branches were labeled their main Y-lineages and their possible related ancestral East Asians. **(C)** Phylogeny of Chinese minority ethnic groups denoted the different lineages. Different colors denoted the different older upper-stream lineages.

### Genetic connections and population stratification among modern and ancient Eurasians

We explored the genetic relationships and population differentiation among 13,777 modern and 1753 ancient Eurasian individuals based on the clustering patterns in the PCA (principal component analysis), MDS (multidimensional scaling analysis) and other population genetic analyses. PCA patterns in the context of Eurasians distinguished ancient western Eurasians from other East Asians (**Fig. S2A-H**). Both harbored different patterns of the dominant Y-lineages and different clustering branches on the phylogenetic tree (**Fig. S2I-J**). The clustering patterns of modern populations were consistent with the geographical divisions and linguistic affinity (**Fig. 2A**). We observed apparent population stratification between northern and southern East Asians and fine-scale substructure among geographically different but linguistically similar groups (**Figs. 2B-D** and **S3A-B**). Most AN/TB-speaking people were separated from others, but other populations possessed much-overlapped clustering positions. The ancient population from the Iron Age Hanben site was clustered closely with AN-speaking people, and northern Chinese ancients were clustered closely with ST-speaking people. We also explored the fine-scale population relationships among Sinitic and TB-speaking people and found Iron Age Hanben people clustered closely with Han Chinese from Guangxi and Taiwan island. However, Yellow River Basin (YRB) farmers clustered separately from other Han Chinese. Clustering patterns among Sinitic and TB people based on Fst matrixes found the differentiated population structure between northern and southern Han Chinese and the genetic differentiation between northern and southern TB people (**Figs. 2D** and **S3A-B**). We also observed substantial genetic differentiation among Altaic-speaking populations, in which Koreanic and Japonic groups each formed individual clades and separated from other Altaic groups, Mongolic and most Tungusic groups clustered together, and Turkic groups showed a close genetic affinity with some Tungusic groups (**Fig. 2B-C**). However, the fine-scale clustering patterns among AA, AN, HM and TK groups from South China and Southeast Asia indicated extensive gene flow events between these populations. The patterns of phylogenetic relationships and HFS further revealed genetic differences between northern and southern Han Chinese and between northern and southern TB-speaking populations, and gene flow events were identified between geographically close populations, such as between AN and southern Han Chinese and between Altaic and northern Han Chinese (**Fig. 2D**).

**Fig. 2.**
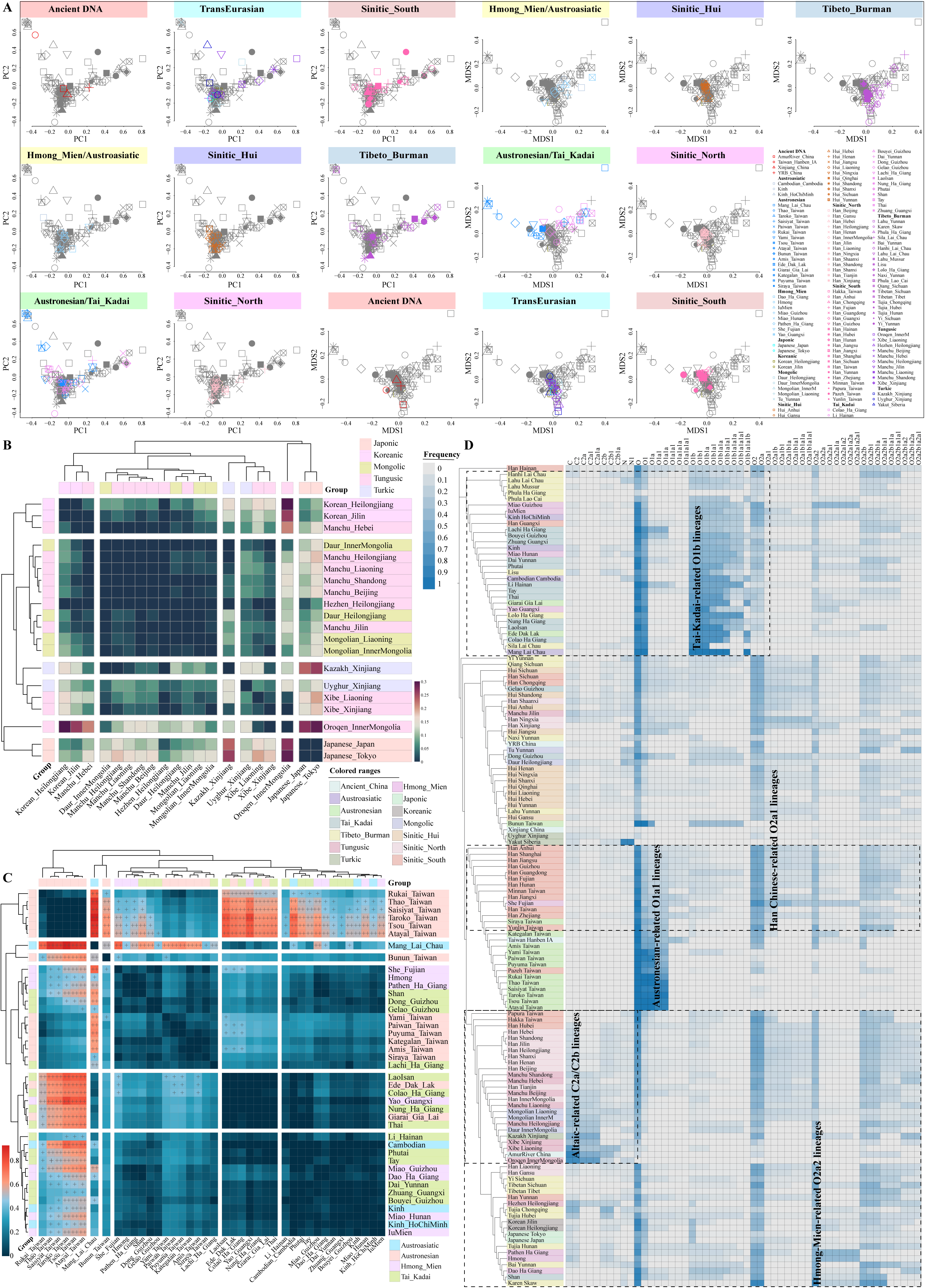
Population genetic structure among 130 modern and four ancient East Asians. **(A)** Principal Component Analysis (PCA) and Multidimensional Scaling plots (MDS) based on the top two components showed the genetic similarities and differences between 134 populations, including 13,886 individuals. **(B-C)** Heatmap showed genetic affinity among Altaic-and Southern Chinese indigenous and Southeast Asian populations. The genetic Fst matrix among Sino-Tibetan-speaking populations was presented in **Fig. S3**. **(D)** Phylogenetic relationships reconstructed based on the Fst matrix and frequency distribution of major Y-chromosome lineages.

We further integrated populations based on their linguistic and ethnic features to explore the genetic affinity among language or ethnicity-based meta-populations via estimating genetic distances and clustering patterns (**Fig. S3C-H**). AN-speaking Saisiyat, Thao, Taroko, Atayal and Tsou from the island of Taiwan clustered together and separated first from other reference populations (**Fig. S3C**). Other distantly separated branches consisted of populations mainly from TK-speaking people, other geographically close AN (Ede and Giarai) and southern TB (Sila and Lolo) speakers. AN branch and TK-dominant cluster were relatively closer to each other than other Asian reference populations, which provided uniparental genetic evidence for the shared or common origin hypothesis of AN and TK language families. Genetic differentiation between populations from these two branches and others was confirmed via Neighbor-joining (NJ)-based phylogenetic relationship, HFS of major founding lineages, and clustering patterns inferred from the PCA and MDS (**Fig. S3D-F**). Fine-scale genetic differences between Sinitic-related Han Chinese and TB people and genetic distinctions among linguistically different populations were visually visualized here. We should highlight that the phenomena of close relationships among most linguistically distinct populations are common here, suggesting massive population movements and gene flow events occurred in the past. Finally, to assess whether paternal lineages provided some evidence supporting the current consensus of language family classification and explore the genetic relationships between linguistically defined meta-populations, we merged all populations based on their linguistic affinities and conducted population genetic analysis based on the genetic distances and haplogroup frequency distributions (**Fig. S3G-H**). We observed a close clustering relationship between TK and AA based on the genetic distances and NJ-related phylogeny. Our language-defined NJ tree also showed a close relationship between Mongolic, Tungusic and ancient ARB, Turkic and ancient Xinjiang people, Koreanic and Japonic, and AN and ancient Hanben (**Fig. S3H**). We provided robust paternal genetic evidence to support the Chinese people’s complex admixture landscape and multifaceted interactions with ancient Eurasians. The simple mergence of geographically distant populations must cause one controversial pattern that may be caused by strategy and sampling biases. We should also pay more attention to the statistically-introduced errors or uncertainties here, which could be gradually overcome in the following WGS-based population genomic studies. We suggested further merging populations by combining the clustering patterns observed in autosomal whole-genome variations and geographical proximity, such as the merged integrated genomic dataset via the administrative divisions.

### The Y-chromosomal diversity landscape shaped by complex population migration and admixture events

Patterns of the observed genetic diversity and paternal genetic structure suggested that complex multiple ancient migration and admixture events may have contributed to the formation of the gene pool of Chinese populations. We subsequently systematically tested how many ancestral sources influenced the paternal genetic composition of Chinese populations and explored the geographical distribution of identified lineages and their correlation with ancient pastoralists and farmers’ Holocene expansion. The haplogroup information of ancient Eurasian and modern East/Southeast Asian populations was collected to comprehensively characterize the patterns of paternal genetic diversity of ethnolinguistically diverse Chinese populations. The final haplogroup dataset contained 115 ethnically or geographically diverse modern Chinese populations, covering 43 officially recognized or unidentified ethnic groups from all provincial-level administrative divisions except Hong Kong and Macau. Moreover, to explore the geographical origin and distribution patterns of dominant paternal lineages in China, we merged all participants into geographically defined meta-populations and estimated the general geographical distribution patterns (**Figs. 3**-**4** and **S4-7**). There were four dominant paternal macro-haplogroups (C-M130, D1-M174, N-M231 and O-M175) in all included Chinese populations, accounting for about 92% of the Chinese Y-chromosomes; the rest consisted of E-M96, F-M89, G-M201, H-L901, I-M170, J-M304, L-M20, Q-M242, R-M207 and T-M184. Finally, we systematically explored how ancient technological innovation and human migrations influenced the Chinese paternal genetic landscape and identified the flowing four ancient migration and admixture events that enriched Chinese diversity (**Fig. 5**).

**Fig. 3.**
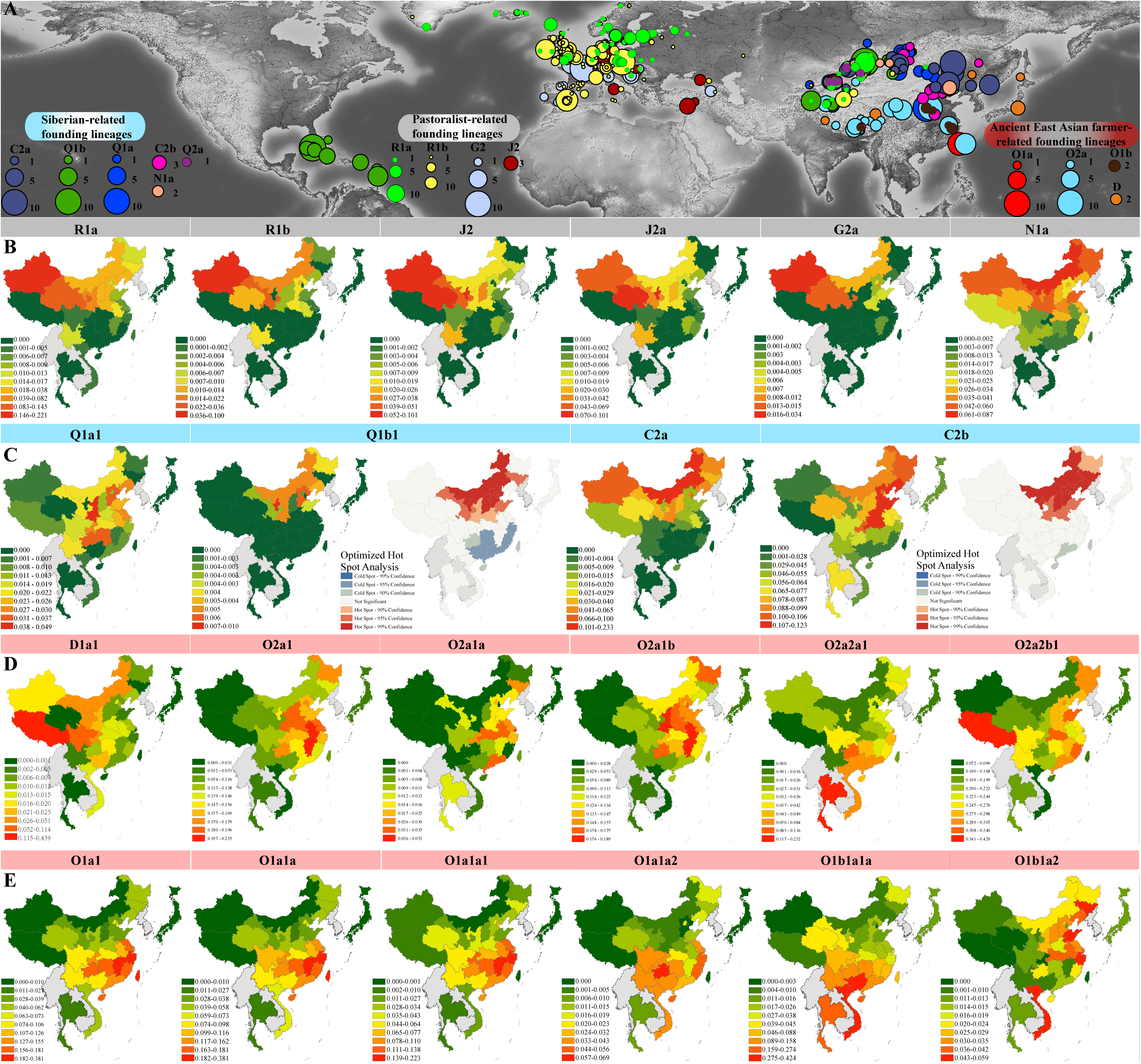
The frequency spectrum of Chinese-dominant Y-chromosome lineages among ancient Eurasians and modern ethnolinguistically diverse East Asians and Southeast Asians. **(A)** The geographical position of 1284 ancient individuals carrying twelve Y-chromosome lineages is interesting in the present work. Different colors of circles indicated different haplogroups, and the circle size denoted the number of interested haplogroups. **(B-C)** Haplogroup frequency of western-origin and Siberian hunter-gatherer-related lineages among eastern Eurasian populations. Optimized hot spot analysis suggested the geographical origin of the focused lineages. **(D-E)** Haplogroup frequency of sublineages derived from the early East Asian-related D, ancient northern East Asian millet farmer-related O2, and ancient southern East Asian rice farmer-related O1. The hot red color suggested the high-frequency or phylogeographical regions of the studied lineages. The frequency distribution of other sublineages was presented in detail in Figs. S4-7.

**Fig. 4.**
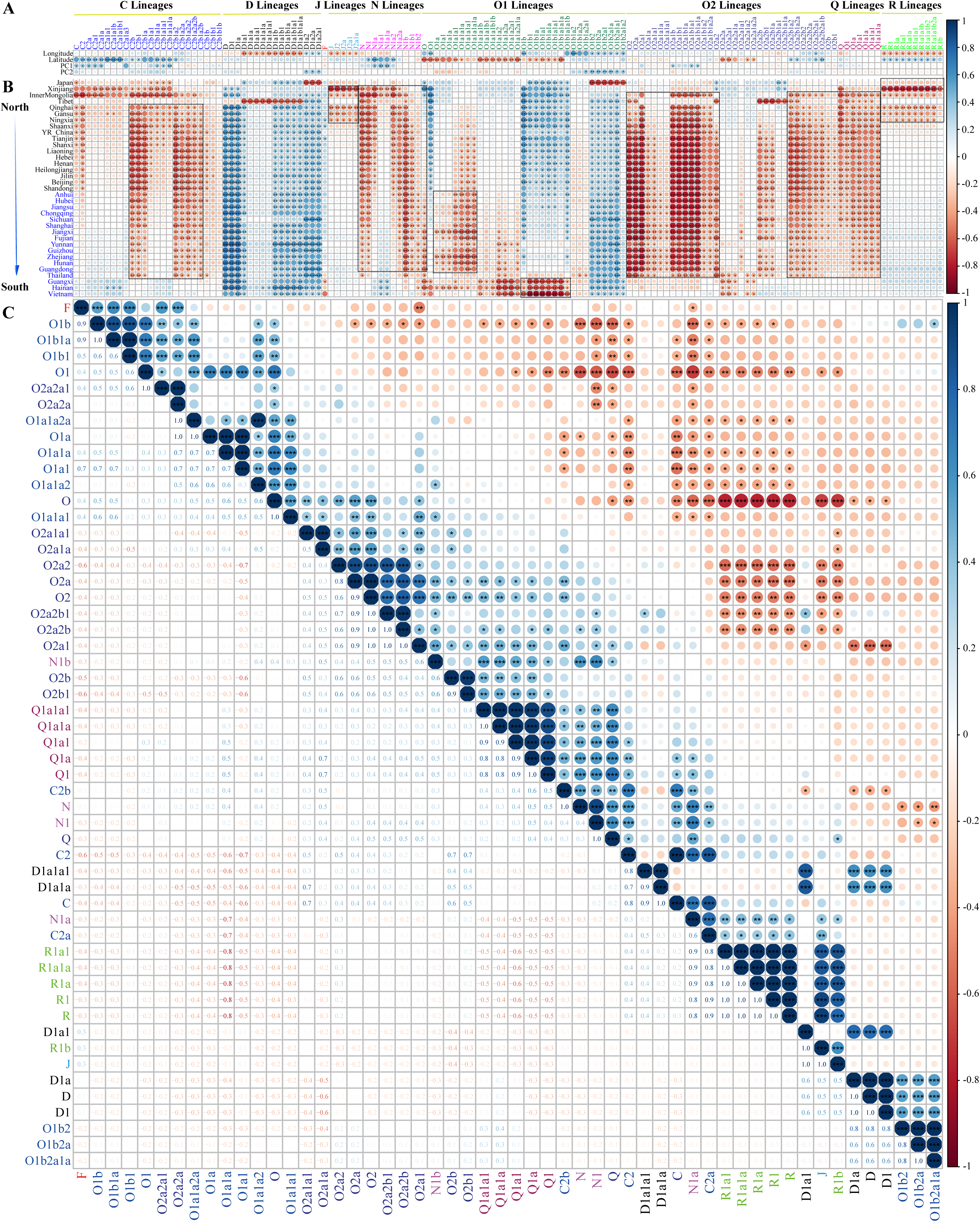
Correlation between frequency of the Chinese-dominant Y-chromosome lineages and other geographical and genetic features. **(A)** The correlation of frequency of Y-chromosome lineages and geographical features of latitude and longitude, as well as with the top two components of the PCA. **(B)** The correlation between the frequency of Y-chromosome lineages and Fst matrix among eastern Eurasian populations. **(C)** The correlation efficient matrix among the pairwise pairs of haplogroup frequency. Asterisk-labeled tested pairs showed that the Spearman correlation is statistically significant. Three asterisks showed p values less than 0.001; two asterisks showed p values ranging from 0.001 to 0.1, and one asterisk indicated the p values ranged from 0.1 to 0.5. The blue color indicated the positive correlation, and the red color indicated the negative correlation.

**Fig. 5.**
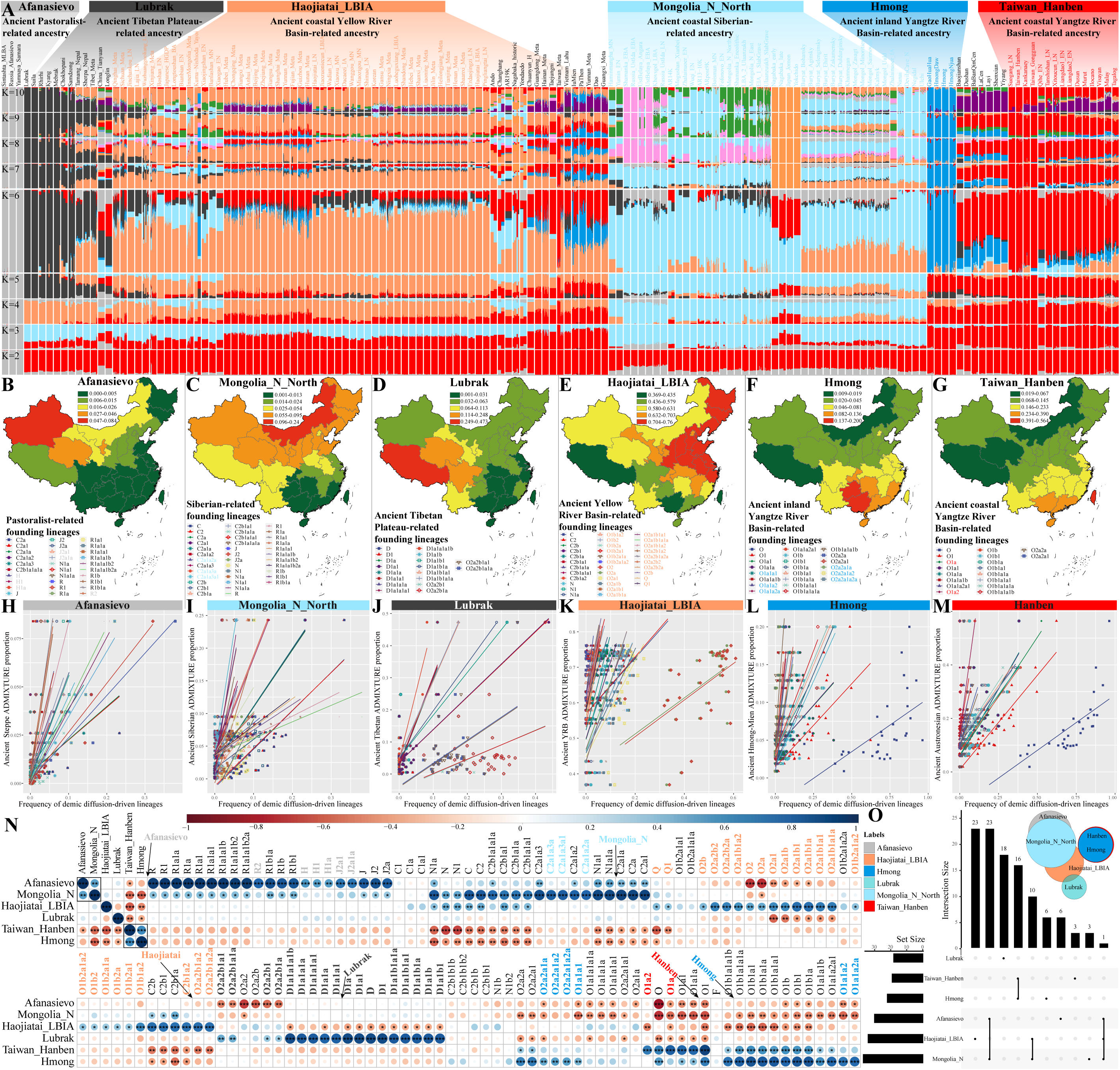
Correlation between autosome-estimated ancestral proportion and Y-chromosome dominant lineages. **(A)** Model-based ADMIXTURE results among modern and ancient East Asians with predefined ancestral sources ranging from 2 to 10. The six-way admixture model with the less cross-validation error was the best-fitted model. **(B-G)** Admixture proportion distribution among different Chinese populations. The red color denoted the highest proportion of one targeted ancestral component. **(H-M)** Scatter plots showed statistically positive correlations between autosome-based ancestral proportion and putatively population-specific founding lineages. **(N)** The correlation between autosome-based ADMIXTURE estimates of ancestral proportion and frequency of Y-chromosome lineages. The correlations between proportions of different ancestral sources were manually removed, and their initial clustering positions were labeled with arrows. **(O)** Venn diagram showed shared and specific lineages among different autosome-based putative ancestral source-related lineages.

### The gene flow from ancient pastoralists and barley farmers’ paternal lineages in Central/South Asia and West Eurasia to East Asia

Historic and prehistoric tans-Eurasian cultural communication events across the southern Bactrian Marianna archaeological complex (BMAC) oasis farming route, the Inner Asian Mountain Corridor (IAMC) biogeography corridor and the northern Yamanaya/Afanasievo steppe pastoralist-related migration route significantly influenced the autosome-related gene pool of ancient people from Altai mountains and surrounding northwestern and northern East Asia (Zhang, Ning, et al. 2021). Whether the human migrations across the Eurasian Steppe and the ancient Silk Road introduced Central/South Asian/West Eurasian-derived paternal lineages to present-day Chinese populations remained unresolved. Our work found that haplogroups R, J, G, Q and their major sublineages had the highest frequency in Northwest China and existed in ancient western Eurasians with high frequency (**Figs. 3A-B** and **S4**). Haplogroup J-M304 most likely evolved in West Asia and was found in significant numbers in present-day West Eurasian populations (Poznik et al. 2016). We observed that most J individuals in China could be subdivided into J2-M172 (especially its sublineage J2a-M410), the distribution center and possible origin of J2a was in the northern Fertile Crescent (**Fig. 3A**) (Grugni et al. 2012), and the current geographical concentration of this lineage largely simulated the agricultural centers (Fuller 2007). Previous findings showed that Northwest Chinese populations, especially Turkic-speaking groups, also carried a strong contingent of J2a (Shou et al. 2010), which was confirmed by our observations. Additionally, ancient individuals belonging to the subclades of J-M304 were identified in the Iron Age Xinjiang populations (Kumar et al. 2022). Chinese individuals carrying G-M201 were also largely restricted to Northwest China. These individuals could be further assigned to G2a (**Fig. 3B**). The spread of G-M201, together with the J2-M172 lineage, was associated with the diffusion of agriculture (**Fig. 3A**) (Semino et al. 2000). Estimates of optimized hot spot analysis (OHSA) confirmed that the diffusion centers of J2a and G2a in China were correlated to Xinjiang and Gansu-Qinghai regions (**Fig. S4A**). Generally, the J/G-derived lineages were likely introduced into China during the eastward migration of Central Asian-related ancestral populations, which may result from the gene flow events mediated by the ancient Silk Road (Zhabagin et al. 2022; He, Yao, et al. 2023).

Haplogroup R-M207 was distributed in ancient western Eurasian and also in modern people in North China at a relatively high frequency, especially in Northwest China (**Fig. 3A-B**), which has been reported to have emerged and diversified in Central Asia and is now common throughout Central/South Asia and Europe (Kayser et al. 2003; Hallast et al. 2021). Only an approximately 24,000-year-old individual (MA1) from the Mal’ta site near Lake Baikal in Siberia belonged to basal haplogroup R (Raghavan et al. 2014). We found that most R individuals could be subdivided into R1-M173, and the haplogroup frequency of its subclade R1a-L146 in China was much higher than that of R1b-M343. In addition, a majority of R1a individuals could be subdivided into R1a1a sublineages, and an extended Y-chromosome investigation revealed R1a1a-M17 as one of the paternal lineages that entered East Asia via the northern route (Zhong et al. 2011). R1b subclades are frequently found in Western Europe (**Fig. 3A**). The estimation of the spatiotemporal distributions of R-M207 subclades supported a West Asian origin of R1b and a subclade carrying the M269 mutation (R1b1a1b-M269) quickly spread to Europe and diffused across Western Europe (**Fig. 3A**) (Myres et al. 2011; Olalde et al. 2018). Haplogroup R2-M479 was found in East Asia at an extremely low frequency, which has been geographically concentrated in Central/South Asia and recently spread from South Asia to North China via the northern route (Zhong et al. 2011; Di Cristofaro et al. 2013). Ancient DNA studies showed that several Mongolia_EIA_Sagly_4 and Mongolia_LBA_MongunTaiga_3 samples that carried more than 40% West Eurasian-related ancestry belonged to R1a1a sublineages (Wang, Yeh, et al. 2021) and several Bronze and Iron Age Xinjiang individuals belonged to R1b sublineages (**Fig. 3A**) (Zhang, Ning, et al. 2021; Kumar et al. 2022). Our findings were consistent with the high frequency of these paternal lineages identified in ancient pastoralists of Yamanaya and Afanasievo populations in the western Eurasian Steppe (Narasimhan et al. 2019; Kumar et al. 2022). The influence of Central/South Asian/West Eurasian-related ancestors could also be reflected by the sporadic occurrence of minor haplogroups, such as E-M96, F-M89, H-L901, I-M170, L-M20 and T-M184, which generally revealed relatively extensive and recent gene flow from these ancestral populations to China.

To further confirm the pastoralist-related population migrations were conferring the reshaped patterns of western Eurasian-related lineages in our studied populations, we estimated the correlation between haplogroup frequency and geographical (longitude and latitude) and genetic features (PC1-2, haplogroup frequency, Fst matrix and autosome-based admixture proportion under the best-fitted models). The frequency of R-related lineages was correlated with latitude and strong genetic affinity with northwestern modern and ancient Chinese populations (**Fig. 4A-B**). R lineages and their sublineages were also observed with a strong correlation (**Fig. 4C**). To illuminate the direct genetic contributions from ancient sources to modern people, we constructed one best-fitted six-source admixture model to explore the ancestral components and proportions of eastern Eurasians and found ancestral proportion decreased gradually from their archeologically attested origin centers (**Fig. 5A-G**). If ancient population movement events directly dispersed the patterns of lineage frequency of ethnolinguistically diverse modern populations, we expected to observe a strong positive correlation between autosome-based admixture proportion of the putative ancestral source and the frequency of the founding lineages. Interestingly, we found a strong correlation between Afanasievo-related ancestry and multiple C, J, N and R sublineages (**Fig. 5H** and 5N). Our results from the HFS of modern and ancient populations, phylogenetic origin inference and multiple factor correlations suggested that western barley and pastoralist people may have promoted the formation of the aforementioned founding lineages.

The dominant Siberian hunter-gatherers’ paternal lineages are widely distributed in China Ancient DNA studies identified an ancestral component that represented the lineage related to Neolithic hunter-gatherers who once lived in the Russian Far East, the Baikal region and Mongolian Plateau, which was referred to as Ancient Northeast Asian (ANA) ancestry (**Fig. 5A** and 5C) (Jeong et al. 2020; Mao et al. 2021). We found that the ANA ancestry made varying contributions to the surrounding spatiotemporally distinct ancient populations, who harbored high proportions of haplogroups Q, R, C and N (Jeong et al. 2020; Wang, Yeh, et al. 2021; He, Yao, et al. 2023). These lineages in the gene pools of modern Mongolic, Tungusic, Turkic and other groups had a significant positive correlation with the proportion of ANA-related ancestry (p values less than 0.05, **Fig. 5N**). Haplogroup Q-M242 was scattered in Chinese populations at extremely low frequencies and its subclades showed different distribution patterns in North and South China (**Fig. 3C**). This lineage might have originated in Central Asia and southern Siberia 15,000Z25,000 years ago. Then, its subclades continued to spread from these regions over the past 10,000 years (Huang, Pamjav, et al. 2018). A major subclade, Q1a1a-M120, was unique to East Asians and found in Han Chinese at a relatively high frequency, which proved to be a crucial lineage of Han Chinese and underwent an in-situ expansion in Northwest China between about 5,000 and 3,000 years ago (Sun et al. 2019). Ancient genomes revealed that most of the Ulaanzuukh_SlabGrave individuals and a Mongolia_LBA_CenterWest_4 individual who carried a minimum proportion of West Eurasian-related ancestry (< 20%) belonged to Q1a1a or its sublineages (**Fig. 3A**) (Wang, Yeh, et al. 2021). Venn-based shared ancestry-correlated lineages also revealed that Q and R lineages were the shared lineages among Yamnaya and ANA-associated lineages (**Fig. 5O**). Besides, one middle Neolithic Yangshao individual and some ∼3,000-year-old Hengbei people from Shanxi belonged to Q1a1a-M120 (Zhao et al. 2014; Ning, Li, et al. 2020), suggesting that ancient individuals carrying Q1a1a contributed to Han Chinese at least 6,000 years ago. Haplogroup Q1b-M346 was rare in China and distributed most intensively in the crossroad between Siberia and North China (**Fig. S4B**), which was reported to have a wide distribution across Asia (Huang, Pamjav, et al. 2018). Some Bronze and early Iron Age individuals from Mongolia were genotyped Q1b1 sublineages (Wang, Yeh, et al. 2021), and some Bronze and Iron Age Xinjiang individuals belonged to Q1b or its subclades (**Fig. 3A**) (Zhang, Ning, et al. 2021; Kumar et al. 2022).

One of China’s most prevalent paternal lineages is haplogroup N-M231, especially its subclade N1-CTS3750. Genetic evidence suggested that N-M231, which presumably originated in South China, spread northward into North China and Siberia during the late Pleistocene-Holocene as the climate warmed and then migrated westward on a counter-clock path from southern Siberia/Inner Asia (Rootsi et al. 2007; Shou et al. 2010; Zhong et al. 2011). Cui et al. conducted Y-chromosome analyses on ancient individuals dating to 6500 to 2700 BP from the West Liao River (WLR) Basin. They found that N-M231 was the leading paternal lineage in Northeast China in the Neolithic Age, and its haplogroup frequency declined gradually over time (Cui et al. 2013). We found that N1a-F1206 showed a relatively high frequency in North China, while N1b-F2930 showed a relatively high frequency in South China, especially in low-altitude Southwest China (**Fig. S4B**). Previous findings provided evidence for the differential distribution patterns of N1a and N1b. That is, the differentiation of the two may have occurred in North China, and then N1a-F1206 migrated northward to areas outside East Asia, while N1b-F2930 migrated southward to South China and became one of the major paternal lineages of TB groups, especially Yi people (Rootsi et al. 2007; Shi et al. 2013; Ilumae et al. 2016; Wang, Song, et al. 2021). N1a1-M46/Tat was a dominating subclade of N1a that probably originated in Northeast Asia (Rootsi et al. 2007; Hu et al. 2015). A sample from the Houtaomuga site in Jilin dating to 7430–7320 years ago belonged to the subclade of N1a1a1a1a-M2117, which was genetically related to early Neolithic ARB individuals, such as DevilsCave_N and Boisman_MN (Ning, Fernandes, et al. 2020). Wang et al. found that several Bronze and Iron Age individuals from the ARB and Mongolia were assigned to N1a or its sublineages, such as a Yankovsky_IA sample (I1202, N1a) and a Mongolia_EIA_SlabGrave_1 sample (I6365, N1a1a1a1a) (Wang, Yeh, et al. 2021). N1a2-F1008/L666 is a dominant paternal type of Uralic populations (Hu et al. 2015), and an ancient DNA study showed that three early Neolithic Shamanka individuals from the Cis-Baikal area were found to belong to N1a2-L666 (de Barros Damgaard et al. 2018). We found that North China and the southwestern part of Northeast China might be the early diffusion center of N1a2 (**Fig. S4B**), which was generally consistent with previous observations (Yu et al. 2023). We observed that N1b-F2930 was mainly distributed in TB-speaking populations in Southwest China and found infrequently in other Chinese populations. However, none of the reported ancient East Asians belonged to haplogroup N1b, and the fine-scale phylogenetic structure of N1b-F2930 needs to be further explored.

Haplogroup C-M130 is a major paternal lineage in East Asia, which might be carried by one of the waves of the earliest settlers, and its diffusion in East Asia was speculated to have begun about 40 thousand years ago (kya) (Ke et al. 2001; Zhong et al. 2010; Wang and Li 2013). C2-M217 was the most widespread subclade and was found with a high frequency in North China (**Figs. 3A**, **3C** and **S4B**). The oldest individual carrying C2-M217 identified so far was AR19K (19,587-19,175 cal BP) in the ARB (Mao et al. 2021). We observed distinct distribution patterns of C2a-L1373 and C2b-F1067, with the highest frequency of C2a-L1373 (sometimes referred to as the “northern branch” of C2-M217) occurring in the Inner Mongolia Autonomous Region and the highest frequency of C2b-F1067 (sometimes referred to as the “southern branch” of C2-M217) in Central, North and Northeast China. We found that most C2a individuals could be further assigned to C2a1a, and its sublineages were often found in Altaic-speaking populations. Specifically, C2a1a1b1-F1756, C2a1a2a-M86 and C2a1a3a-F3796 were the major subclades of C2a1a individuals in China (**Fig. S4B**). The phylogeny reconstruction of C2a1a1b1-F1756 suggested that its two major subclades might be related to the early expansions of Mongolic/Tungusic-related ancestors (Wei et al. 2017). Several studies have shown that C2a1a2a sublineages were widespread in Altaic groups in East Asia and North Asia (Chen et al. 2011; Liu, Ma, et al. 2021; Wang, He, et al. 2021), and some ancestral Tungusic groups in the ARB migrated to the Mongolian Plateau and contributed the genetic components of C2a1a2a to present-day Mongolic/Turkic speakers (Liu, Ma, et al. 2021). C2a1a3a-F3796, also known as the C2*-Star Cluster (C2*-ST), is one of the founding paternal lineages of Mongolic-speaking populations (Zerjal et al. 2003; Wei et al. 2018; Wang, He, et al. 2021). The highly revised phylogenetic tree of C2*-ST and the estimated time of TMRCA of this paternal lineage and its sublineages suggested that the dispersal patterns of C2*-ST correlated with the expansion of Mongolic-speaking populations, whose origins could be traced back to either ordinary Mongolian tribes or an ancient Niru’un Mongol clan (Wei et al. 2018). Wang et al. found that several ancient individuals from the ARB and Mongolia belonged to C2a1a or its sublineages, such as Mongolia_North_N and Boisman_MN (**Fig. 3A**) (Wang, Yeh, et al. 2021), indicating that ANA-related populations contributed significantly to present-day Mongolic, Tungusic and some Turkic groups. We also identified some individuals belonging to C2a sublineages in central and southern Chinese populations, indicating that Proto-Mongolian populations from North Asia migrated southward and spread the C2a sublineages, which was mainly driven by the expansion of the Mongol Empire (Wei et al. 2017; Zhang, Wu, et al. 2018; Li et al. 2020).

The phylogenetic analysis of C2b-F1067 suggested that ancient populations carrying various C2b sublineages contributed significantly to the gene pool of modern Eastern Eurasians (Wu et al. 2020). Our observations showed that the Inner Mongolian Plateau and Northeast China might be the initial dispersal center of C2b (**Fig. S4B**), consistent with a previous finding that North China might be the diffusion center of C2b before 11 kya (Wu et al. 2020). We revealed different geographical distribution patterns of C2b sublineages (**Fig. S4B**). For example, C2b1a1-CTS2657 was distributed in North and Northeast China at relatively high frequencies; C2b1a2-F3880 was prevalent in Northeast, North (except the Inner Mongolia Autonomous Region and Shanxi) and East (mainly Shandong, Jiangsu and Shanghai) China; while C2b1b-F845 has distributed in Central and Southwest (mainly Guizhou) China as well as Southeast Asia at the highest frequency. Significantly, our findings were partially inconsistent with a previous study (Wu et al. 2020), which may be caused by the bias of sampling and reference populations. Our study and several previous studies confirmed the southern origin of C2b1b-F845, and an ancient DNA study further suggested that the origin of this paternal lineage was associated with southern farming populations, who contributed the genetic components of C2b1b to ancient nomadic groups on the Mongolian Plateau (Li et al. 2020; Wu et al. 2020). Statistically significant negative correlations between western Eurasian and Siberian-related lineages suggested that their high frequency in northern East Asians contributed to their genetic differentiation from southern East Asians (**Fig. 4B**). Generally, genetic analyses based on the modern and ancient East Asians suggested the solid genetic connection between Neolithic Mongolian hunter-gatherers and modern East Asians.

Early Asian paternal lineages left traces mainly in the Tibetan Plateau and surrounding regions The ancient genetic connection between Andamanese, Jomon-related indigenous Japanese, and highland Tibetans was evidenced via the shared Palaeolithic autosomal ancestry components and uniparental D lineage (**Fig. 3A** and 3D). We explored the phylogeographical origins of D subclades and found that D1-M174, as one of the four major Y-chromosome haplogroups in East Asians, did occur at high frequency in our YHC (**Fig. S5**), and this lineage had been intensively studied and showed clear southern origin (Shi et al. 2008; Zhong et al. 2011; Qi et al. 2013). Haplogroup D1a exhibited high proportions and clustering centers in the Tibetan Plateau (TP) and the Japanese archipelago (**Fig. S5**). Significantly, most D1a individuals (> 95%) could be subdivided into D1a1a-M15 or D1a1b-P99. We observed that D1a1a sublineages were found frequently among TB-speaking populations in Southwest China and at low frequency in other Chinese populations, whereas D1a1b sublineages occurred at the highest frequency on the TP. Previous studies showed that D1a1a-M15 diverged from D1-M174 during the migration of D1-M174 to East Asia, and then D1a1a-M15 migrated northward through western Sichuan to Gansu-Qinghai region and might have entered the Himalayan area along the Tibetan-Yi corridor; D1a1b-P99, primarily its subclade D1a1b1-P47, diverged from D1-M174 on the TP, which resulted in D1a1a sublineages being widely distributed in China and D1a1b sublineages being specific to Tibetan populations (Qi et al. 2013; Wang et al. 2014). Sublineages of D1a were uniquely maximized in Tibetan populations, which was confirmed by the genetic contribution from North China millet farmers to highlanders via the revised Y-chromosome phylogeny, correlation with O lineages and Lubrak-related TP ancestry (**Fig. 4C** and 5D). The estimated gene flow events, Lubrak-related D sublineages and the close genetic connections between North China and TP highlanders contributed to Chinese people’s genetic diversity patterns. Interestingly, the frequency of four lineages (O2a2b1, O2a2b1a, O2a2b1a1 and O2a2b1a1a) also strongly correlated with the Lubrak-related ancestry further confirmed the Neolithic expansion from YRB promoted the peopling of the highland TP (**Fig. 5J**).

### Ancient northern East Asian millet farmer-related farming-language-people southward dispersal

Archeological and historical documents suggest that YRB was the cradle of ancient Chinese civilization. Ancient DNA of millet farmers from Houli, Yangshou and Longshan cultures showed that all ST people originated from North China (Wang, Yeh, et al. 2021). Wang et al. investigated the genetic profiles of spatiotemporally different Guangxi populations and identified southward genetic influence related to Shandong ancients (Wang, Wang, et al. 2021). Yang et al. identified persistent southward gene flow from Shandong to the coastal southeastern East Asia (Yang et al. 2020). We also identified Haojiatai-related ancestry dominant in Chinese populations, strongly correlated with the O, Q, C and N lineages. Haplogroup O-M175 is primarily found in East and Southeast Asians and has two main subclades: O1-F265 and O2-M122. The origin, diversification and expansion of these two lineages and their sublineages were suggested to be associated with the spread of millet and rice farmers from the domestication agriculture centers of the YRB and Yangtze River Basin (YZRB) (**Fig. 5K-M**). However, the extent to which ancient northern East Asians reshaped the paternal genetic diversity of modern southern East Asians remained to be comprehensively assessed. Thus, we evaluated the geographical distribution of O-related lineages and explored the correlation with the estimated millet farmer ancestry (**Fig. 5K**). Ancient O2 lineages were widely distributed in northern China and the TP (**Fig. 3A**). Our findings showed that haplogroup O2-M122, especially its subclade O2a-M324, is one of the major paternal lineages in East Asian populations and is also highly prevalent in Southeast Asian populations (**Fig. 3D** and **S6**) and a strong correlation within different subclades (**Fig.4C**). Y-chromosome evidence based on 15 Y-SNPs demonstrated a southern origin of O2-M122 and suggested that its northward migration in East Asia occurred ∼25Z30 kya (Shi et al. 2005). We observed that O2a-M324 was widely distributed in China’s coastal and surrounding areas with extremely high frequency (**Fig. S6B**), indicating that the ancestors carrying O2a-M324 were likely to migrate along the coast and then spread to other East Asian areas and Southeast Asia. Yan et al. sequenced 3.9 Mbp NRY region of 78 East Asians and identified three star-like Neolithic expansions [Oα (O2a2b1a1-M117), Oβ (O2a2b1a2a1a-F46) and Oγ (O2a1b1a1a1a-F11)] in haplogroup O2a-M324, which suggested that the male-mediated expansion in China mainly occurred during the Neolithic Age (Yan et al. 2014). Ancient DNA evidence revealed that a middle Neolithic individual from the WLR belonging to Hongshan culture was assigned to haplogroup O2a-M324 (Ning, Li, et al. 2020). Yu and Li reviewed the origin of Chinese ethnic groups, language families and civilizations from the perspective of Y-chromosome and revealed that O2a-M324 originated in northeastern China and was associated with the development of Hongshan culture. We did observe that the highest frequency of haplogroup O2a-M324 occurred in northeastern China’s Heilongjiang (**Fig. S6B**). However, its highest haplogroup frequency was also observed in some eastern coastal provinces, such as Shandong, Shanghai, Fujian and Guangdong, which may be caused by the bias in included ethnically diverse populations and sample sizes. Additionally, OHSA results indicated that the middle and lower YRB could be the early diffusion center of O2a-M324.

We observed distinct distribution patterns of O2a1-L127.1 and O2a2-JST021354/P201 (**Fig. S6**). O2a1 occurred at the highest frequency in East China, and its haplogroup frequencies decreased in the surrounding areas (**Fig. S6C**). At the same time, O2a2a-M188 was distributed in Southeast Asia at relatively high frequencies and then spread from south to north in East Asia with a decrease in the haplogroup frequency. O2a2b-P164 was prevalent in China and occurred at the highest frequency in the TP. We found that the overwhelming majority of O2a1 individuals could be further subdivided into O2a1b-JST002611, and it was widely distributed in different Chinese ethnic groups, especially in Han populations (**Fig. S6D**), which was consistent with previous genetic findings (Wang et al. 2013; Yao et al. 2017). The low-frequency distribution of O2a1b and its sublineages in TB-speaking populations suggested that this lineage did not play a significant role in forming TB speakers. There are two main sublineages in O2a1b defined by F11 (O2a1b1a1a1a) and F238 (O2a1b1a2a), respectively (Wang et al. 2013; Yao et al. 2017). We observed that O2a1b1a1a1a-F11 was more frequently distributed in China than O2a1b1a2a-F238, especially in geographically diverse Han populations. Haplogroup O2a1b1a1a1a was distributed in East China at higher frequencies, which was consistent with previous findings that O2a1b1a1a1a-F11 experienced a rapid expansion probably in eastern Han Chinese about five kya (Wang et al. 2013). Wang et al. found that the F238 mutation was likely to occur in Proto-Han-Chinese about seven kya, but it was difficult to clarify the origin of this mutation based on the Y-STR haplotype diversities of O2a1b1a2a-F238 (Wang et al. 2013), which was confirmed by the observation in this study that haplogroup O2a1b1a2a was mainly restricted to Northeast, North and East China (**Fig. S6E**). We found that the initial diffusion center of O2a1b sublineages was likely to be the middle and lower YRB (**Fig. S6D**). Zhang et al. reported genome-wide data from 12 ancient samples dating to 6500 to 2500 years ago in China and identified an O2a1b1a1a1a-F11 sample from the Banpo site, indicating that the Yangshao people may have induced the generation of haplogroup O2a1b1a1a1a (Zhang, Lei, et al. 2018). Ning et al. found that a late Neolithic individual from the Erdaojingzi site belonging to the Lower Xiajiadian culture in the WLR (WLR_LN) could be assigned to haplogroup O2a1b-JST002611 and WLR_LN derived their ancestry mainly from YRB-related ancestral populations (88 or 74%) (Ning, Li, et al. 2020).

Most O2a2a subclades were distributed with high frequencies in South China and Southeast Asia (**Fig. S6F-G**). Previous studies suggested that O2a2a1a2-M7 was one of the founding lineages of HM-speaking populations (Xia et al. 2019; Kutanan et al. 2020; Liu, Xie, et al. 2021) and found at a high frequency in the Daxi people in the middle reaches of the YZRB (Li et al. 2007), demonstrating that the Daxi people might be the ancestors of present-day HM speakers. We found that O2a2b sublineages were widely distributed in China. O2a2b1-M134 was one of the main subclades of O2a2b, which was found frequently among ST-speaking populations, and the highest frequency occurred in Tibetan populations (**Fig. S6H**). Two O2a2b1 subclades (O2a2b1a1-M117 and O2a2b1a2a1a-F46) showed star-like expansions (Yan et al. 2014). The highest frequency of O2a2b1a1 was observed in the TP and Southeast China, but the OHSA results showed that the early diffusion center of O2a2b1a1 might be the TP (**Fig. S6H**). We found that O2a2b1a2, the upstream haplogroup of O2a2b1a2a1a, was distributed in Northeast, North and East China at higher frequencies than other regions (**Fig. S6H**). Ancient DNA studies revealed that O2a2b1 sublineages were identified in ancient individuals from the Shimao site belonging to the Longshan culture (Shimao_LN), from the Jinchankou, Lajia and Mogou sites belonging to the Qijia culture (Upper_YRB_LN) and from the Dacaozi site (Upper_YRB_IA) (Li et al. 2017; Ning, Li, et al. 2020). Wang et al. demonstrated that O2-M117-F5 (called Oα-F5) was one of the founding paternal lineages of modern TB groups and the Yangshao people in the middle YRB carrying Oα-F5 migrated southwestward to the TP at about six kya and mixed with the local D-M174 populations to form the first layer of the genetic profiles of modern TB-speaking populations (Wang et al. 2018). Furthermore, several genome-wide SNP studies revealed that Core Tibetans (Tibetans from the TP) had more than 70% ancestry related to Upper_YRB_LN or more than 85% ancestry related to YRB_MN (Wanggou_MN and Xiaowu_MN) belonging to the Yangshao culture (He, Wang, Zou, et al. 2021; Wang, Yeh, et al. 2021). Therefore, the development of millet agriculture, migration of millet farmers and admixture with geographically diverse indigenous populations resulted in the current distribution patterns of diverse O2a-M324 sublineages, with the leading subclades of O2a1b-JST002611 and O2a2b1-M134.

### Ancient southern East Asian rice farmer-related founding lineages from the YZRB left a massive genetic legacy in China and Southeast Asia

South China was another agricultural origin center of rice domestication and was proposed as the hometown of HM, TK, AA and AN people. The best-fitted ADMIXTURE models revealed inland Hmong- and coastal Hanben-related ancestral components widely distributed in southern Chinese populations and correlated with most O1 subclades (**Fig. 5F-G** and **5L-O**). Recent ancient DNA also illuminated that ancient YZRB rice farmers genetically reshaped the patterns of ancient YRB millet farmers and Southeast Asian modern and ancient populations (Yang et al. 2020; Wang, Wang, et al. 2021). We here explored the phylogeographical features of O1 sublineages to illuminate the impact of paleo-genomically attested gene flow events on the paternal genetic diversity of Chinese and Southeast Asians. We found that O1-F265 was found at high frequencies in South China, Southeast Asia and the Japanese archipelago, and its subclade O1a-M119 was frequently found in southeastern Chinese populations, while O1b-M268 was frequently distributed in southwestern Chinese and Southeast Asian populations (**Fig. S7**). O1a sublineages were mainly distributed in AN-, TK- and Sinitic-speaking populations in Southeast Asia and South China, indicating that AN and TK speakers shared a common patrilineal ancestor and extensive gene flow existed between Han Chinese and AN/TK-related ancestral populations. Their genetic interaction and admixture signatures were also identified via the autosome-based admixture models (Chen et al. 2022; Wang, He, et al. 2022; Liu et al. 2023). We observed that O1a1a1 and its sublineages (except O1a1a1b) were found at high frequencies in Southeast China, and their early dispersal centers might be the middle and lower reaches of the YZRB and the southeast coast (**Fig. S7B**). Previous research showed that the paternal lineages observed in the Liangzhu populations from the Yangtze River Delta were mostly O1a-M119 and in the Taiwan Hanben individuals were mostly O1a1 sublineages (Li et al. 2007; Wang, Yeh, et al. 2021), and rice farmers carrying O1a-M119 in the YZRB were likely to be the direct ancestors of modern TK and AN speakers, they migrated southward along the southeast coast of China to Southeast/Southwest China and mainland Southeast Asia (Liu et al. 2020; Wang, Yeh, et al. 2021; Wang, Wang, et al. 2021; Wang, He, et al. 2022). We observed that O1a1a1b was found at the highest frequency in Hainan, especially in Li people, and its haplogroup frequency showed a general south-to-north decline, suggesting that Li-related ancestors made significant contributions to other southern Chinese populations (**Fig. S7B**). Our observations showed that O1a1a2 and its sublineages were found at high frequencies in Southwest China and Vietnam, and Southwest China was likely to be the initial diffusion center of these paternal lineages (**Fig. S7C**). However, the high-resolution phylogeny of O1a-M119 revealed that O1a1a2-F4084+/K644-were found at higher frequencies in the Yangtze River Delta, while its southwestern sublineage O1a1a2a1a-K644, a founding paternal lineage of TK-speaking populations, showed an apparent south-north dispersal trend starting from Hainan Island (Sun et al. 2021). Substantial differences in the coverage of target populations and genotyping methods may lead to partial inconsistencies in the results of different studies. Generally, the migrations of rice farmer-related ancestral populations largely contributed to the observed distribution patterns of O1a sublineages.

We observed that O1b-M268 was distributed in Southwest China, Southeast Asia and the Japanese archipelago at high frequencies, which could be subdivided into three major subclades (O1b1a1-PK4, O1b1a2-Page59 and O1b2-P49) that showed significantly different distribution patterns (**Fig. S7D**). O1b1a1 and its sublineages were mainly distributed in Southwest China and Southeast Asia, and a significant sublineage O1b1a1a-M95 mainly existed in AA groups in these regions and was also an important paternal lineage of TK-speaking populations (Zhang et al. 2014; Kutanan et al. 2019; Song et al. 2019; Macholdt et al. 2020). However, the geographical origin and migratory routes of O1b1a1a remain controversial. Ancient DNA evidence revealed that the Wucheng people in Jiangsu along the YZRB and two ancient individuals from the Hengbei site in Shanxi approximately 3,000 years ago carried the O1b1a1a-M95 lineage (Li et al. 2007; Zhao et al. 2014). We observed that O1b1a2 and its sublineages were relatively rare in East Asia and mainly distributed in East China, the southwestern part of Northeast China and Vietnam, especially in Han Chinese (**Fig. S7E**), consistent with previous findings (Yan et al. 2011). Ancient genomes from North China revealed that a middle Neolithic individual from the Wanggou site belonging to the Yangshao culture was assigned to O1b1a2-Page59 (Ning, Li, et al. 2020). Haplogroup O1b2-P49 was found at the highest frequency in Japan, followed by Northeast China, but the detailed phylogenetic structure of this lineage remains to be further reconstructed (**Fig. S7F**). Our identified patterns of genetic diversity from O1 lineages suggested that ancient rice farmers from South China significantly influenced the gene pool of populations from South China and Southeast Asia. In general, complex population movements and admixture events contributed to the formation of modern and ancient East Asians. To illuminate the origins of different Chinese-dominant subclades and the demographic processes of ethnically/geographically diverse modern Chinese populations, we should design a systematic sampling strategy, conduct whole Y-chromosome sequencing and fully retrieve spatiotemporally distinct ancient individuals for more comprehensive analyses.

## Conclusion

Genetic evidence from autosome-based ancient DNA has revolutionized our understanding of human population genetic history; however, ancient genetic legacy inferred from the ancient Y-chromosome was limited, as the lower copy number than mitochondrial genomes. We launched the YanHuang cohort to capture the Y-chromosome diversity of ethno-linguistically diverse Chinese populations via genotyping 919 individuals from 39 ethnic minority groups using our recently developed high-resolution YHSeqY3000 panel and merged with the Y-chromosome genomic database of 14,611 people, including 1753 ancient people, to explore the formation process of ancient Chinese Y-chromosome genetic diversity landscape. Ancient DNA data collected from ancient Eurasian populations and modern integrated data included 115 ethnolinguistically or geographically distinct modern Chinese populations belonging to 47 officially recognized or unidentified ethnic groups and covered all provincial-level administrative divisions except Hong Kong and Macau. Our results identified multiple founding lineages related to ancient western herder Euraisan, Siberian Fisher-Hunter-Gatherer and millet and rice farmers from Yellow and Yangtze River Basians contributed to the patterns of geography-related population paternal genetic stratification. We illuminated the strong correlation between the frequency of subsistence-model-related founding lineages and the autosome-based admixture proportion of putative ancestral sources, latitude and differentiated north-to-south genetic matrix, suggesting ancient population movements and extensive admixture between incomers and indigenous populations was the major mechanism for the formation of the evolutionary spectrum of the Y-chromosome landscape. We emphasized the importance of combining high-deep whole-genome sequencing data of modern and spatiotemporally different populations to validate and further characterize the paternal evolutionary history of East Asians.

### Materials and Methods Study participants

To fully characterize the panorama of Y-chromosomal diversity in China, we collected saliva samples from 919 participants from 39 ethnolinguistic groups belonging to Sinitic (Hui), TB (Bai, Derung, Lahu, Lisu, Lhoba, Nakhi, Pumi, Qiang, Tibetan, Tujia and Yi), TK (Bouyei, Dai, Dong, Gelao, Li, Maonan, Shui, Zhuang and Mulao), HM (Miao, She and Yao), AN (Gaoshan), AA (Jing and Wa), Mongolic (Daur, Dongxiang and Mongolian), Tungusic (Hezhen, Manchu and Xibe), Turkic (Kazakh, Kyrgyz, Uzbek and Uyghur), Koreanic (Korean) and IE (Russian) language families (Table S1). The sampled individuals were descendants of self-identified members of the given ethnic groups and their grandparents lived in the sampling districts for at least three generations. This study was approved by the Medical Ethics Committee of West China Hospital of Sichuan University (2023-306) and conducted in accordance with the Helsinki Declaration of 2013 (Jama 2013). In addition, we obtained informed consent from each participant.

### DNA extraction, sequencing and genotyping

We extracted genomic DNA using the QIAamp DNA Mini Kit (QIAGEN, Germany). DNA concentrations were quantified using the Qubit dsDNA HS Assay Kit based on the standard protocol on a Qubit 3.0 fluorometer (Thermo Fisher Scientific). The Y-specific target sequences with 50X coverage were generated on the Illumina platform (Illumina, San Diego, CA, USA) using the custom-designed YHSeqY3000 panel. The raw sequencing reads were mapped to the human reference genome GRCh37 using BWA v.0.7.13 (Li and Durbin 2009).

### Haplogroup classification and phylogeny construction

We first conducted NRY haplogroup classification using in-house scripts. The NRY haplogroups were also classified using HaploGrouper (Jagadeesan et al. 2021) and Y-LineageTracker (Chen et al. 2021) based on the Y-DNA Haplogroup Tree 2019Z2020 (https://isogg.org/tree/index.html), respectively. For the complete dataset of 919 samples, a maximum-likelihood phylogenetic tree was constructed via MEGA X (Kumar et al. 2018), which was then visualized using iTOL (Letunic and Bork 2021). The haplotype and haplogroup diversity were estimated using the following formula: 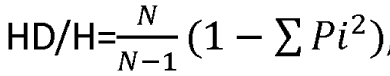, where N denotes the total number of observed haplotypes or N-1 haplogroups and Pi denotes the frequency of the i-th haplotype or haplogroup. We only extracted populations with a sample size greater than 30 for the estimation of HD and H values. Haplogroup frequency spectrum estimation and clustering analysis

### Dataset composition

We incorporated previously published haplogroup information of 11,979 East Asian individuals from 79 populations retrieved from previous fragmentation studies (Trejaut et al. 2014; Lang et al. 2019; Song et al. 2019; Xie et al. 2019; Wang, Song, et al. 2021; Wang, He, et al. 2021; Song et al. 2022; Wang, Song, et al. 2022), the 1KGP and the Human Genome Diversity Project (HGDP) (Poznik et al. 2016; Bergstrom et al. 2020), 879 individuals from 28 Southeast Asian populations (Kutanan et al. 2019; Kutanan et al. 2020; Macholdt et al. 2020), 131 ancient East Asians from Xinjiang (Xinjiang_China), ARB (AmurRiver_China), YRB (YRB_China) and South China (Hanben_IA) (Jeong et al. 2020; Wang, Yeh, et al. 2021; Zhang, Ning, et al. 2021; Kumar et al. 2022) and 1622 ancient western Eurasians from Allen Ancient DNA Resource into this study to estimate the HFS and depict the population structure of ethnolinguistically diverse Chinese ethnic groups. A total of 13,777 present-day individuals were collected from 12 linguistically different groups, covering 22 provinces, five autonomous regions and four municipalities in China as well as Thailand and Vietnam, including 135 AA-, 693 AN-, 285 HM-, 75 Japonic-, 35 Koreanic-, 994 Mongolic-, 863 TK-, 1338 TB-, 260 Tungusic-, 291 Turkic-, 1 IE-speaking individuals (excluded from population genetic analysis), 805 Sinitic-speaking Hui, 3248 northern Han Chinese and 4754 southern Han Chinese (Tables S1 and S3). The retrieved haplogroups were manually revised based on the variant information and the Y-DNA Haplogroup Tree 2019Z2020. In order to estimate the spatial distributions of different paternal lineages more conveniently, we integrated haplogroup data to generate meta-populations based on the geographical region, ethnicity and language family. We estimated haplogroup frequency based on different levels of terminal haplogroups. We conducted population genetic analysis based on individual populations with a sample size over ten or meta-populations with a sample size larger than 30. Haplogroup frequency of all upstream lineages was used to explore corresponding deep population genetic history.

### Population structure inference

We calculated pairwise Fst genetic distances based on the HFS using Y-LineageTracker (Chen et al. 2021). The NJ phylogenetic tree was constructed using MEGA (Kumar et al. 2016) and modified using the iTOL online tool. MDS was conducted based on the genetic distance matrix using the cmdscale R tool (https://itol.embl.de/itol.cgi), and PCA was conducted based on the HFS using Y-LineageTracker (Chen et al. 2021).

### Spatial statistics correlated the phylogeographical origin of founding lineages

The haplogroup frequency of one province-defined population at different levels of terminal haplogroup was calculated using Y-LineageTracker with level parameters ranging from 0 to 6. Geographically diverse Chinese populations were merged based on the administrative distinctions of provinces, and populations from the island and mainland of Southeast Asia were integrated based on the country. Using ArcMap, we investigated the geographical distribution patterns of dominant haplogroups in Chinese populations by OHSA (Getis-Ord General G) and spatial autocorrelation analysis (Moran’s I). The clusters (hot and cold spots) revealed by OHSA roughly mirrored the possible geographical origin or diffusion center of a specific haplogroup and its mirror regions showed the general distribution pattern of that haplogroup.

### Phylogenetic relationship reconstruction

Fasata data was used to reconstruct the phylogenetic tree using using Y-LineageTracker (Chen et al. 2021). Network relationship of shared haplotype was explored via the popart (Leigh et al. 2015) software.

### Autosome-based ADMIXTURE estimation

We collected 445 ancient individuals from 88 Eurasian populations and 1325 geographically different modern individuals from 62 populations from our merged 10K_CPGDP database to form the autosome-based dataset, including ancient Chinese populations from inland and coastal southern and northern East Asia, hunter-gatherers from Mongolia and western Yamnaya-related pastoralists. We modeled the admixture proportions of geographically different populations via best-fitted ADMIXTURE. We pruned the autosome-based dataset using PLINK (Chang et al. 2015) with the parameters of “--indep-pairwise 200 25 0.4” and “--allow-no-sex” and then ran ADMIXTURE with the predefined ancestral sources ranging from 2 to 15 (Alexander et al. 2009). We used cross-validation error values to identify the best-fitted admixture models and used the admixture proportion of modern populations to correlate autosome-based ADMIXTURE and haplogroup frequency.

### Correlation between lineage frequency and ADMIXTURE-based ancestry proportion

We first calculated the haplogroup frequency of geographically defined meta-populations. Chinese populations were grouped based on the provincial administrative region. We cut all tested lineages at the nine levels and identified 139 common lineages with a frequency larger than 0.05 in at least one population, 177 low-frequency lineages and 165 rare lineages. We then explored the Person correlation between haplogroup frequency and longitude, latitude and their inter-correlation and statistical significance using corrplot R packages. Followingly, we merged all Chinese populations as one super-population and then defined all common lineages with a frequency larger than 0.01 or 0.05. We also used the corrplot R package to test the correlation between ADMIXTURE proportion and haplogroup frequency.

Declarations

### Ethics approval and consent to participate

The Medical Ethics Committee of West China Hospital of Sichuan University approved this study. This study was conducted following the principles of the Helsinki Declaration.

### Consent for publication

Not applicable.

### Availability of data and materials

All haplogroup information was submitted in the supplementary materials. We followed the regulations of the Ministry of Science and Technology of the People’s Republic of China. The raw genotype data required controlled access. Further requests for access to raw data can be directed to Guanglin He and Mengge Wang.

### Funding

This work was supported by grants from the National Natural Science Foundation of China (82202078).

### Authors’ contributions

G.H., M.W. B.Z. and C.L. conceived and supervised the project. G.H. and M.W. collected the samples. K.L., K.Z., Y.H., G.H. and M.W. extracted the genomic DNA and performed the genome sequencing. G.H., M.W. and K.L. did variant calling. M.W., Y.H., K.L., H.Y., Z.W., S.D., L.W., H.Y., Q.S., J.Z., R.T., J.C., Y.S., X.L., C.W., H.S., Q.Y., L.H., L.Y., J.Y., S.N., Y.C., J.Y., K.Z., B.Z., C.L., G.H. performed population genetic analysis. G.H. and M.W. drafted the manuscript. G.H., M.W., B.Z. and C.L. revised the manuscript.

## Acknowledgments

We thank all volunteers who participated in this project.

